# Downstream branches of receptor tyrosine kinase signaling act interdependently to shape the face

**DOI:** 10.1101/2024.12.10.627829

**Authors:** Nicholas Hanne, Diane Hu, Marta Vidal-García, Charlie Allen, M. Bilal Shakir, Wei Liu, Benedikt Hallgrímsson, Ralph Marcucio

**Author notes:** co-first authors (Nicholas Hanne, Diane Hu). Grant Sponsor and Number: CIHR Foundation grant 159920, NIH NIDCR R21-DE028198, NIH NIDCR F32-DE030359, National Science and Engineering Research Council (NSERC), #238992-2; Canada First Research Excellence Fund 2022-00015, Canadian Foundation for Innovation #36262.

## Abstract

**Background:** Previously we found that increasing fibroblast growth factor (FGF) signaling in the neural crest cells within the frontonasal process (FNP) of the chicken embryo caused dysmorphology that was correlated with reduced proliferation, disrupted cellular orientation, and lower MAPK activation but no change in PLCy and PI3K activation. This suggests RTK signaling may drive craniofacial morphogenesis through specific downstream effectors that affect cellular activities. In this study we inhibited three downstream branches of RTK signaling to determine their role in regulating cellular activities and how these changes affect morphogenesis of the FNP.

**Results:** Small molecule inhibitors of MEK1/2, PI3K, and PLCy were delivered individually and in tandem to the right FNP of chicken embryos. All treatments caused asymmetric proximodistal truncation on the treated side and a mild expansion on the untreated side compared to DMSO control treated FNPs. Inhibiting each pathway caused similar decreased proliferation and disrupted cellular orientation, but did not affect apoptosis.

**Conclusions:** Since RTK signaling is a ubiquitous and tightly regulated biochemical system we conclude that the downstream pathways are robust to developmental perturbation through redundant signaling systems.

**Bullet points:** Inhibiting three downstream effectors of receptor tyrosine kinase (RTK) signaling (MEK1/2, PLCy, and PI3K) in the frontonasal process of chicken embryos caused similar mild truncation of growth. Combining all three inhibitors had a slightly stronger effect on truncation.

Individual inhibitors did not have specific effects on cellular proliferation, apoptosis, or cellular orientation.

The downstream branches of RTK signaling likely have shared interdependent effects on cellular activities that contribute to morphogenesis.

## Introduction

Tissues of the head develop from primordia that are comprised of neural crest cells, surface ectoderm and mesoderm. These primordia evaginate and fuse to form the structures of the upper jaw and palate.^1,2^ Alterations to morphogenesis of the facial primordia can cause asymmetry, prevent apposition and fusion of primordia, or cause premature fusion of primordia.^3,4^ The process of fusion is complex and controlled by regional differences in cellular proliferation, apoptosis, and differentiation, and also directed cellular migration and oriented cellular behavior.^5,6^ Signaling via receptor tyrosine kinases (RTKs), such as fibroblast growth factor receptors (FGFRs) and platelet-derived growth factor receptors (PDGFRs), are required for normal formation of the face.^7–13^ Alterations to these pathways can lead to malformations including facial asymmetry. RTK signaling directs cellular proliferation, apoptosis, differentiation, migration, and oriented cell behaviors through several downstream intracellular signaling pathways.^14–16^

Previously, we found that activating the fibroblast growth factor (FGF) pathway in neural crest cells of chick embryos, using replication competent avian sarcoma retroviruses (RCAS) encoding either *Fgf8*, or *FgfR2^C278F^*, a constitutively activated receptor, slowed proximodistal expansion of the frontonasal process (FNP) and widened the midface.^11^ Increasing fibroblast growth factor (FGF) signaling decreased cellular proliferation, which was correlated with proximodistal hypoplasia as well as increased expression of FGF pathway antagonists. Additionally, the more severely malformed embryos exhibited more random orientation of the Golgi apparatus in relation to the nuclei,^11^ suggesting that polarized cell behaviors may be involved in facial morphogenesis as in the limb bud.^17–19^ The RCAS::*Fgf8* virus decreased the amount of phosphorylated ERK1/2, a signaling molecule downstream from FGFR, which has been shown to stimulate cellular proliferation and direct Golgi orientation.^11,20^ In the mesenchyme, activation of the FGF pathway increased expression of FGFR inhibitors *Spry1*, *Spry2*, *Spred2*, and *Dusp6,* which could explain decreased activation of ERK1/2, but the amount of phosphorylation of PLCγ and PI3K, two other signaling pathways downstream from FGFR, were unaffected.^11^ FGF signaling alters several cellular functions simultaneously that appear to be specific to individual downstream signaling pathways and likely drive craniofacial morphogenesis. Here, we use small molecules to specifically inhibit the MEK1/2, PI3K, and PLCγ pathways, major downstream effectors of RTK signaling, to determine their role in regulating cellular activities and how these changes affect morphogenesis of the FNP.

## Results

Beads containing small molecule inhibitors (U0126 to inhibit MEK1/2, LY294002 to inhibit PI3K, and U73122 to inhibit PLCγ) were implanted into the lateral edge of the globular process of the FNP of chicken embryos at stage HH19/20, a period when the FNP is expanding rapidly but has not yet fused with the maxillary process, to determine the contribution of individual pathways downstream of RTK activation on facial morphogenesis (Figure 1). Shape change and cellular activities were quantified 24 hours later at stage HH24/25 (Figure 1).

**Figure 1:**
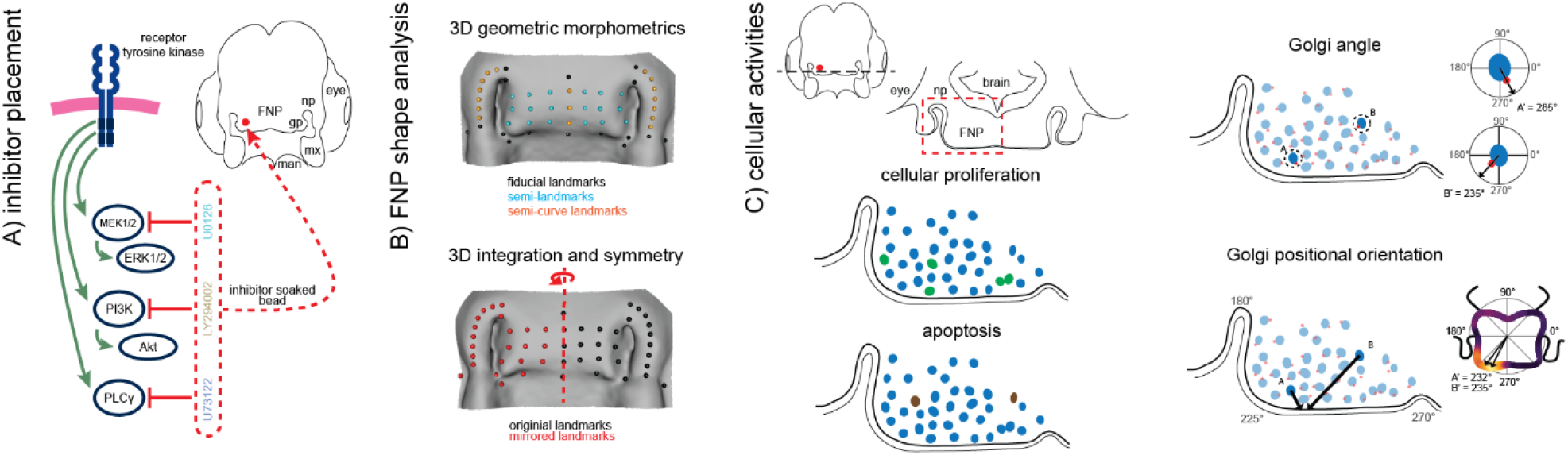
Experimental overview. A) Beads soaked in small molecule inhibitors were implanted into the right side of the chicken frontonasal process at stage HH19/20. Immunohistochemistry labelling of phosphorylated forms of ERK1/2, Akt, and PLCγ confirmed pathway inhibition and specificity of the inhibitors 6 and 24 hours after implantation. B) Shape change was quantified 24 hours after treatment at stage HH24/25 using geometric morphometric analysis. Landmarks were placed on 3D reconstructions of uCT scans. B top) Geometric morphometric analysis was performed on all landmarks to determine the effect of inhibitor treatment on overall shape change, and the symmetric and asymmetric components of shape. B bottom) The effect of each treatment on the contralateral (untreated) and treated sides was further analyzed by making new faces composed of one side of the face mirrored over the opposite side. C) Proliferation, apoptosis, and Golgi orientation were measured in transverse sections of the FNP, adjacent to the bead, 24 hours after treatment. Golgi orientation was analyzed with two methods. Golgi angle (top) was calculated as the angle between a nucleus and its neighboring Golgi body. The Golgi positional orientation (bottom) was calculated as the position along the tissue section where the Golgi body is pointing towards.

### Inhibiting downstream effectors of RTK activation, individually or in concert, caused truncation of the facial primordia and increased variance in morphogenesis

Each of the three inhibitors caused similar mediolateral and proximodistal truncation of the FNP after 24 and 72 hours (Figure 2A and 2B). Qualitatively, we observed a variable effect of any individual inhibitor treatment on FNP shape – many treated samples had obvious truncation of the FNP while others resembled DMSO control samples. We quantified FNP shape change 24 hours after treatment using generalized Procrustes analysis (GPA) on three-dimensional landmark data (Figure 1B) and visualized shape differences with principal component analysis (PCA) and canonical variates analysis (CVA). Shape was significantly associated with facial size, quantified as the centroid size of the landmark coordinates (p = 0.001). The relationship between shape and size varied significantly by treatment group (p = 0.028). Despite this interaction effect, there were no significant pairwise differences between treatment groups and DMSO treated embryos (DMSO vs any treatment p > 0.48) or between all inhibitor treated embryos compared to DMSO (p = 0.761). The first principal component for shape variation captures symmetric expansion of the midline of the FNP, which likely reflects the allometric relationship between shape and size. To remove the allometric variation we regressed the residuals of the first principal component out of the original landmark coordinates and re-ran the GPA with the new coordinates (Fig 2C). All subsequent analysis of shape was performed with these transformed data.

**Figure 2:**
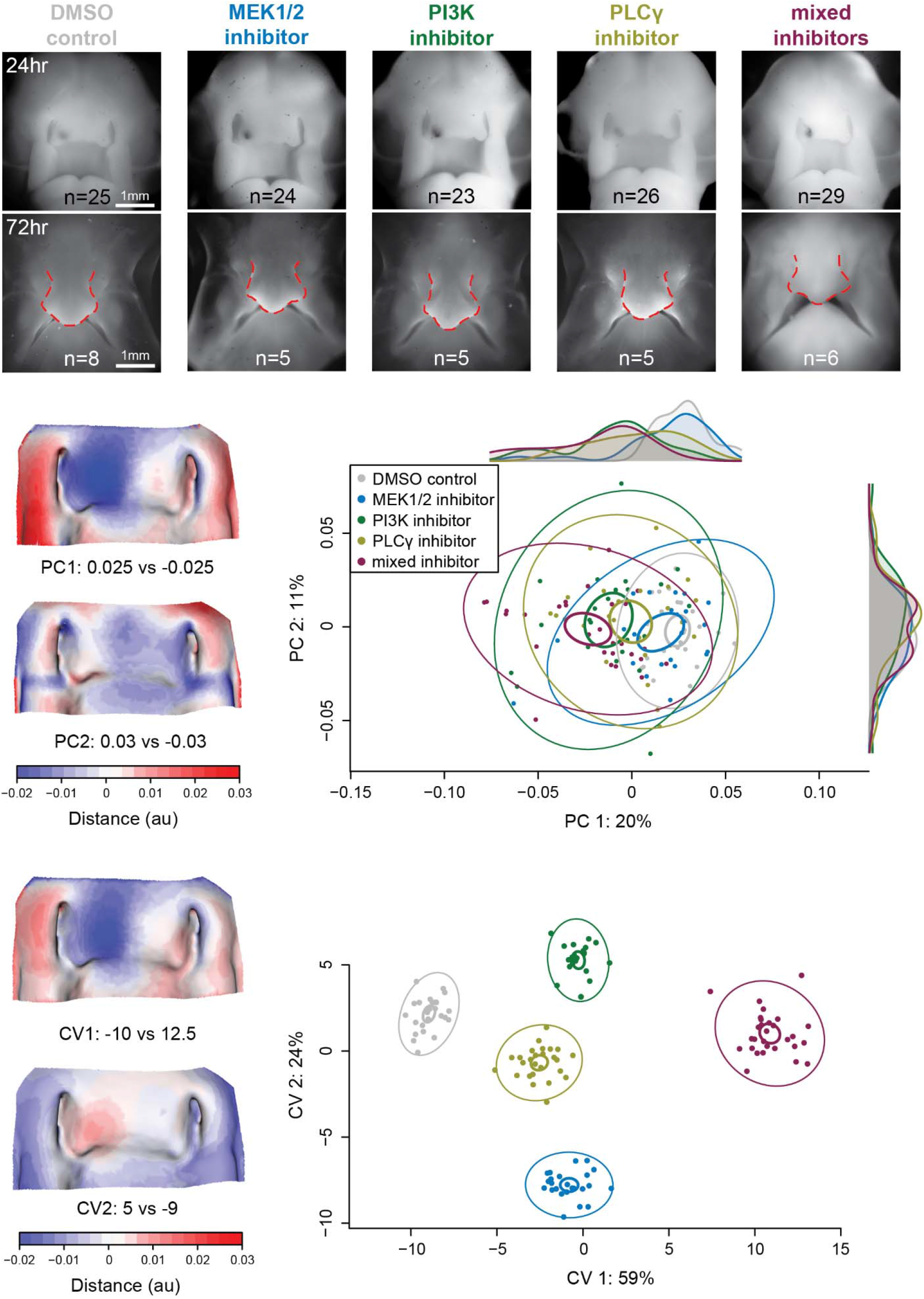
Quantitative and qualitative shape change after downstream RTK pathway inhibition. Inhibiting the downstream targets of RTK activation – MEK1/2 with U0126, PI3K with LY294002, PLCγ with U73122, and all three inhibitors combined – caused truncation of the FNP A) 24 hours (stage HH24/25) after bead placement and B) 72 hours (stage HH28) after bead placement. The red outlines are added to demonstrate the asymmetry present in the treated side of the face after 72 hours. C) 3D geometric morphometric analysis revealed significant effects on FNP shape and increased variance in FNP shape 24 hours after treatment compared to DMSO treated controls. The heatmaps represent differences in shape within the first two principal components from a shape near the DMSO mean shape to an extreme shape near the edge of the range of observed treated embryos. The first PC heatmap shows that the inhibitors caused asymmetric proximodistal truncation on the treated side compared to DMSO. The second PC heatmap shows subtle variation in the amount of truncation on the contralateral, untreated side of the FNP. D) Canonical variates analysis separated DMSO from individual inhibitors from the combined inhibitors along the first axis. Ellipses on the principal component plot and canonical variates plot represent 95% confidence intervals of standard deviation (outer, thinner ellipses) and 95% confidence intervals of standard error of the mean (inner, thicker ellipses).

In a principal components analysis of the size-regressed data, the first PC (20% of variance) captures variation that separates the inhibitor treated FNPs from the DMSO treated FNPs. This involved proximodistal truncation on the treated side of the FNP compared to the contralateral side (Figure 2C). The second principal component contains subtle variation in the amount of truncation of the globular process on the untreated side and midline (Figure 2C). This variation is likely due to placement of the bead. These same shape differences separated treatment groups in the first two components of canonical variate analysis, which represent the two largest sources of variation between groups that vary the least within each group (Fig. 2D). Treatment had a significant effect on shape of the FNP (p = 0.001), and most inhibitor treatments caused significant shape change compared to the DMSO control treated samples (p = 0.060 for inhibiting MEK1/2; p = 0.001 for inhibiting PI3K; p = 0.002 for inhibiting PLCγ; Figure 2C). Inhibiting PLCγ had similar effect on shape of the FNP as inhibiting MEK1/2 or PI3K (p = 0.072, and p = 0.158, respectively); all other individual treatment comparisons were significantly different (p = 0.001). Treatment also had a significant effect on FNP shape variance (p = 0.001) and each inhibitor treatment tended to increase variance compared to DMSO control treatment (p = 0.093 for inhibiting MEK1/2; p = 0.003 for inhibiting PI3K; p = 0.091 for inhibiting PLCγ; Figure 2C). The effect of each inhibitor on shape variance was similar (p > 0.05 for all inhibitor vs inhibitor treatment comparisons).

We determined whether the downstream pathways were affecting FNP shape through shared or independent mechanisms by simultaneously inhibiting MEK1/2, PI3K, and PLCγ with beads containing a cocktail of U0126, LY294002, and U73122, respectively. The ‘mixed inhibitor’ treatment had a significantly different effect on mean FNP shape than most individual inhibitor treatments (p = 0.001 compared to DMSO; p = 0.001 compared to MEK1/2; p = 0.056 compared to PI3K; p = 0.002 compared to PLCγ) but had similar effect on variance of FNP shape (p = 0.066 compared to DMSO; p > 0.05 for all mixed inhibitor vs individual inhibitor treatment comparisons) (Figure 2C). Although mixed treatment effect on mean shape was significantly different from most individual inhibitor treatment, the mixed inhibitor treatment did not have as strong an effect as we have observed previously when inhibiting FGFR directly with SU5402, suggesting these and other downstream pathways have shared, inter-dependent effects on the cellular processes that drive morphogenesis.^21^

Inhibiting downstream effectors of RTK activation caused both symmetric and asymmetric shape change despite only treating the right side of the FNP (Figure 3A and 3B). Treatment had a significant effect on symmetric shape change (p = 0.001) that was driven primarily by variation in proximodistal expansion of the lateral nasal processes and the FNP midline. Embryos separated from DMSO in the first principal component have wider lateral nasal processes, but this may be due to having relatively truncated FNPs due to the inhibitors. Inhibiting MEK1/2 had similar effect on symmetric shape change as DMSO treatment and inhibiting PLCγ (p = 0.075 and p = 0.067 respectively), while inhibiting PLCγ had similar effect as inhibiting PI3K (p = 0.329). All other pairwise comparisons were significantly different (p < 0.05, Figure 3A).

**Figure 3:**
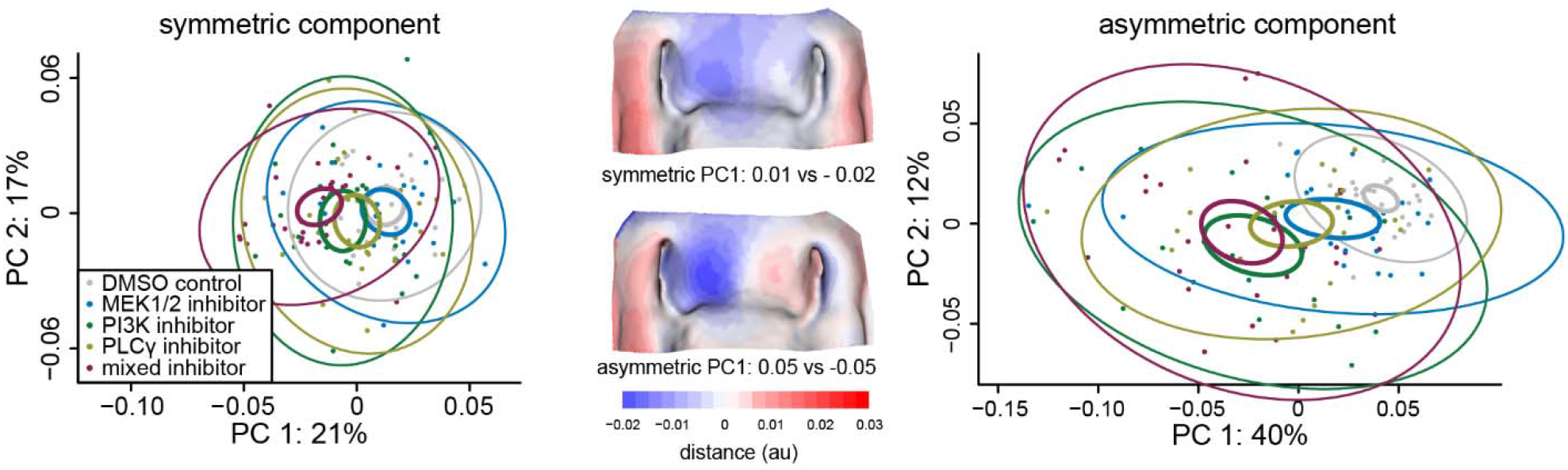
Inhibitors caused A) symmetric and B) asymmetric shape change despite unilateral treatment. Aside from the MEK1/2 inhibitor treatment, all treatments were significantly different from DMSO control. Ellipses represent 95% confidence intervals of standard deviation (outer, thinner ellipses) and 95% confidence intervals of standard error of the mean (inner, thicker ellipses).

Treatment also significantly increased symmetric shape variance (p = 0.001, Figure 3A), but only inhibiting PI3K significantly increased variance compared to DMSO (p = 0.006). Each inhibitor treatment had a significant effect on asymmetric shape change that was driven by proximodistal truncation on the treated side (p = 0.001) and increased asymmetric shape variance (p = 0.001, Figure 3B). Most inhibitor treatments caused significant asymmetric shape change compared to the DMSO control treatment (p = 0.064 for inhibiting MEK1/2; p = 0.001 for inhibiting PI3K; p = 0.003 for inhibiting PLCγ; p = 0.001 for mixed inhibitor, Figure 3B). Inhibiting MEK1/2 caused less asymmetric shape change than inhibiting PLCγ (p = 0.002) and the mixed inhibitor treatment (p = 0.001), but all other treatments had similar effects on asymmetry (p > 0.05, Figure 3B). Each inhibitor treatment increased variance of asymmetric shape change compared to DMSO (p = 0.015 for inhibiting MEK1/2; p = 0.001 for inhibiting PI3K; p = 0.007 for inhibiting PLCγ; p = 0.001 for mixed inhibitor, Figure 3B), but all of the treatments had similar variance in asymmetry compared to one another (p > 0.05 for all inhibitor-inhibitor pairwise comparisons).

We explored the shape change on the untreated side of the FNP more closely by examining covariation between the two sides of the FNP. We used paired two-block partial least squares analysis to quantify how changes to the inhibitor treated side of the FNP covary with the contralateral side. All inhibitor treated FNPs and DMSO control treated FNPs exhibited strong covariation between their treated and contralateral sides with partial least squares analysis (p <= 0.004 for all groups, Figure 4A). The effect size of the two-block partial least square analyses from inhibitor treated FNPs and DMSO control treated FNPs were similar (p = 0.999 for inhibiting MEK1/2; p = 0. 719 for inhibiting PI3K; p = 0.380 for inhibiting PLCγ; and p = 0.221 for inhibiting all three with the mixed treatment), suggesting the inhibitor treatments did not disrupt covariation across sides of the face despite causing asymmetric truncation.

**Figure 4:**
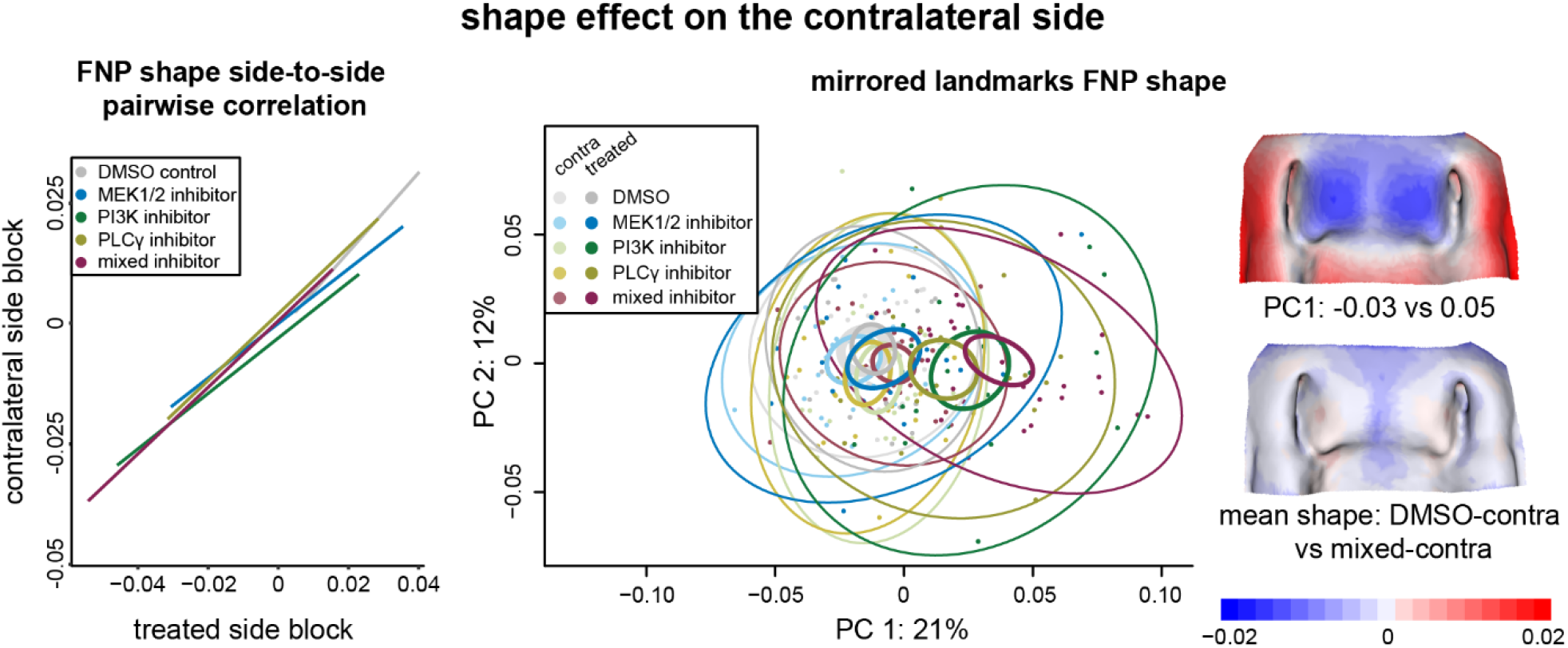
The shape of the contralateral side of the FNP was affected by unilateral inhibitor treatment and continued to covary with the treated side. A) Variation in shape on the treated side of the FNP are strongly correlated with variation in shape on the contralateral side in all treatment groups despite the strong asymmetric shape change observed. The effect sizes of variation between the two sides of the face were not different between any inhibitor treatment and DMSO, suggesting inhibitors did not disrupt covariation between the halves of the FNP. B) Principal components analysis of shape of mirrored FNPs separated contralateral-only FNPs from treated-only FNPs along the first PC. Unilateral inhibitor treatment affected the contralateral side of the face – there were no contralateral-only faces from inhibitor treated embryos that resembled the DMSO-contralateral faces. The ellipses on the principal component plot represent 95% confidence intervals of standard deviation (outer, thinner ellipses) and 95% confidence intervals of standard error of the mean (inner, thicker ellipses).

Finally, we performed geometric morphometric analyses on mirrored FNPs to determine the effect of treatment on the contralateral side and further examine the effect of the inhibitors on the contralateral, untreated side. We mirrored landmarks from the treated or contralateral sides of each FNP to create faces composed of only one side-treatment combination (i.e. ‘DMSO-treated’ FNP is made up of landmarks from the bead treated side and those same landmarks mirrored across the midline instead of the original contralateral side landmarks) (Figure 1B). The mean shapes of the mirrored FNPs separated along the first principal component (Figure 4B) and tended to separate contralateral-only FNPs from treated-only FNPs based on the severity of truncation. No contralateral-only FNPs from inhibitor treated embryos resembled the DMSO-contralateral, but DMSO-contralateral was similar to MEK1/2-treated (p = 0.121). The difference between inhibitor treated contralateral-only FNPs and DMSO-contralateral FNPs was driven by truncation of the midline and expansion of the globular process (Figure 4B heatmap). Similarly, only DMSO and MEK1/2 treated FNPs had treated and contralateral sides that resembled their other halves (p = 0.878 for DMSO-contralateral vs DMSO-treated; p = 0.157 for MEK1/2- contralateral vs MEK1/2-treated). These results further illustrate the extent to which inhibitor treatment caused shape change on the contralateral, untreated side of the face.

### Small molecule inhibitors decreased proliferation but did not affect apoptosis

Inhibiting PI3K decreased proliferation relative to the contralateral untreated side (p = 0.014, n = 6) but inhibiting PLCγ had a weak effect on relative proliferation (p = 0.062, n = 20) and inhibiting MEK1/2 had no effect (p = 0.157, n = 16) (Figure 5). These results are surprising as MEK1/2 is a strong mitogen yet had no effect on proliferation.^22,23^ However, the effects of inhibiting MEK1/2 and PI3K may have been strong enough to decrease proliferation on the treated and contralateral side – proliferation on the contralateral side of MEK1/2 and PI3K inhibited FNPs were lower than the treated and contralateral sides of PLCγ inhibited FNPs (p < 0.001 and p = 0.016, dotted line in Figure 5). None of the treatments caused noticeable differences in cell death compared to DMSO control treated FNPs.

**Figure 5:**
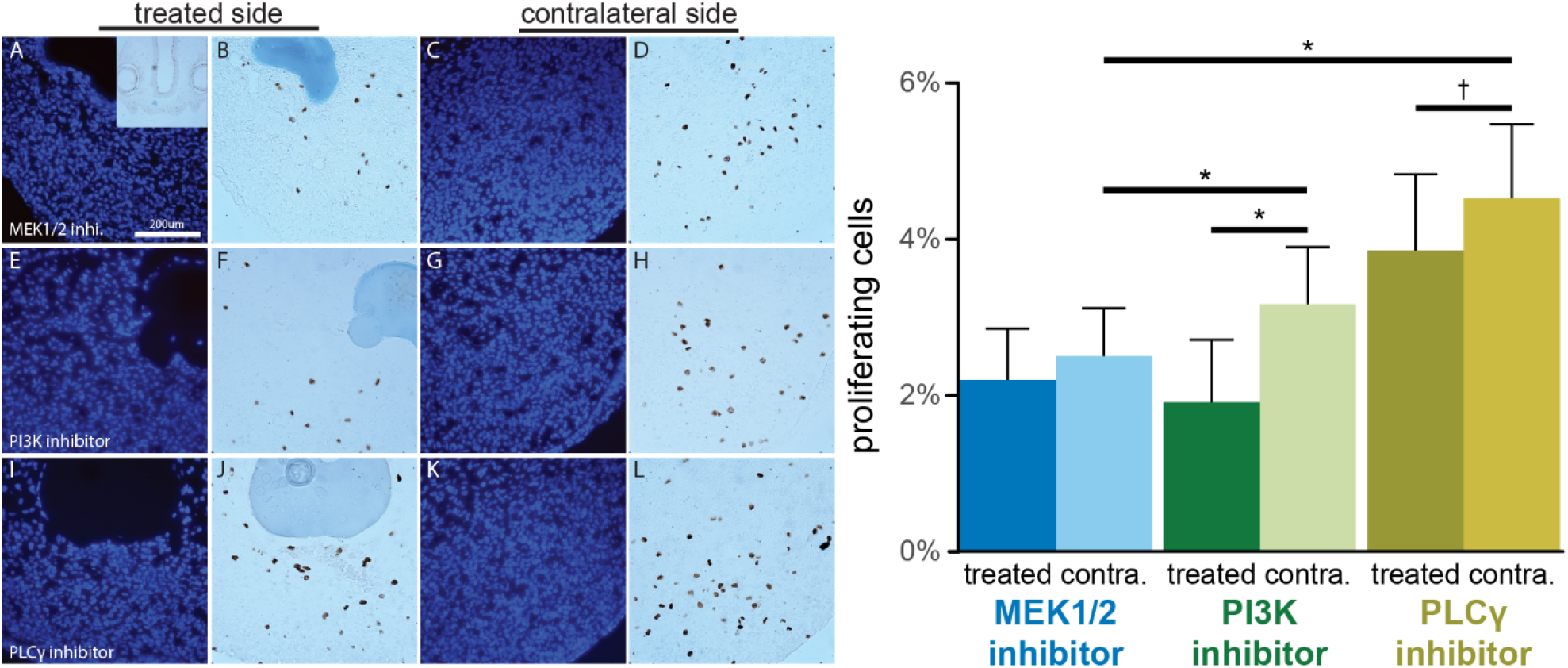
FGF pathway inhibition tended to decrease cell proliferation. Nuclei and proliferating cells were labelled with Hoecsht (first and third image column) and anti-pHH3 antibodies (second and fourth image column). Bargraph) Inhibiting PI3K decreased proliferation on the treated side compared to the contralateral side. MEK1/2 inhibitor and PI3K inhibitor treatment reduced proliferation on the contralateral side compared to the contralateral side of FNPs treated PLCγ inhibitors. * p <0.05; dagger p < 0.1.

### Inhibiting downstream branches of RTK activation disrupted cellular orientation

As cells migrate the Golgi body tends to be located between the nucleus and the direction of growth.^24^ We quantified cellular orientation in the mesenchyme by measuring the angle formed between a cell’s nucleus and its Golgi body (‘Golgi angle’, Figure 1C) and by measuring where along the tissue the Golgi is pointing towards (‘positional orientation’, Figure 1C). Roughly 1,000 nucleus-Golgi body pairs were analyzed in each side for each sample (over 50,000 pairs total). Golgi angle was highly variable in all FNPs (Figure 6 and Table I), with large standard deviations and small mean resultant lengths (1-variance). In the DMSO control FNPs the Golgi are more oriented toward the distal and lateral direction of the ipsilateral side of the FNP, while the individual inhibitor treated samples appear more randomly oriented (Figure 6). We compared each treated side to their contralateral side using the two-sample Watson’s U^2^ test, which tests if two distributions of circular data could have been sampled from the same population. Inhibiting all three downstream pathways with the mixed inhibitor treatment altered Golgi angle on the bead treated side compared to the contralateral side, while inhibiting MEK1/2 and the DMSO control did not (Table I). Inhibiting PI3K and PLCy had a weak effect on Golgi angle compared to the contralateral side. However, each of the inhibitor treatments disrupted orientation compared to a distribution made up of the combined DMSO treated and contralateral sides (Table I, white distributions in Figure 6).

**Figure 6:**
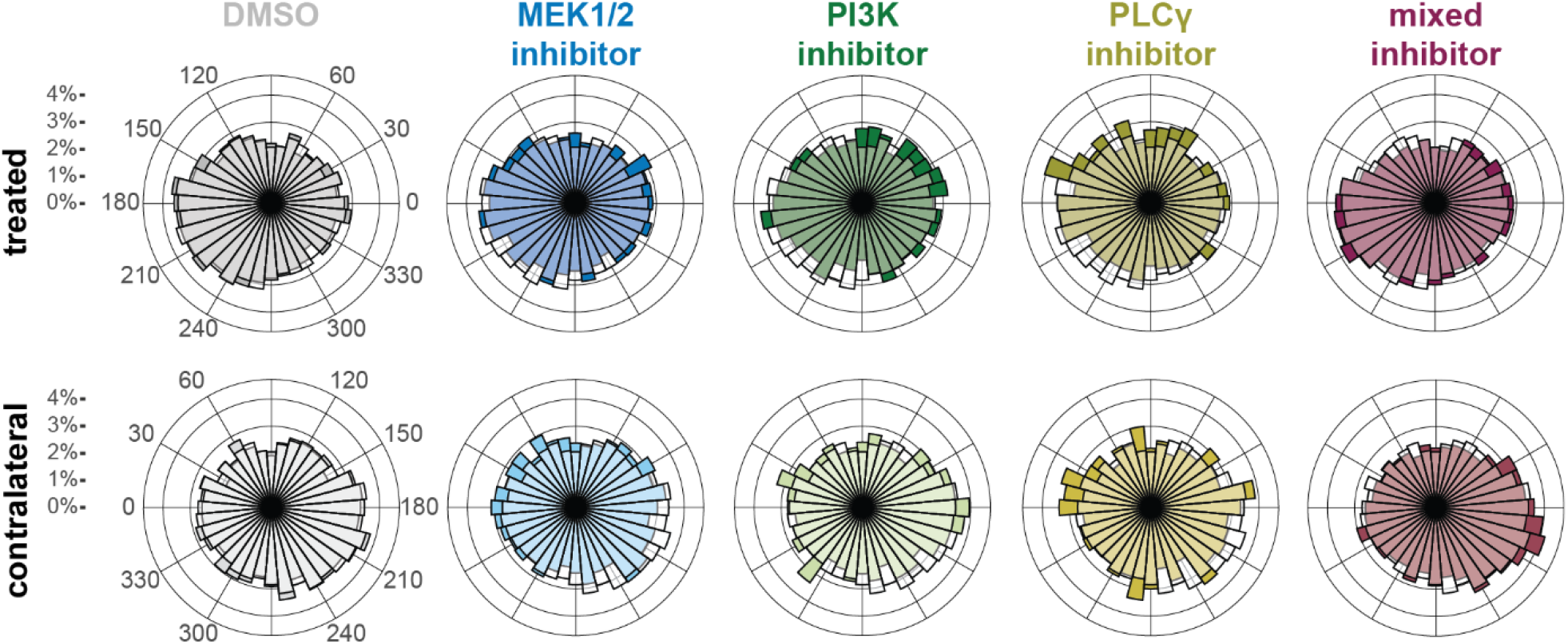
All inhibitor treatments disrupted Golgi angle compared to DMSO control. Angle between a nucleus and Golgi body is illustrated with wind rose histograms of percent of nuclei for each treatment and side combination. Each windrose histogram shows distribution of Golgi-nuclei angle for the treatment-side combination in color as well as the DMSO combined sides distribution in white.

**Table I:** Golgi body angle circular distribution statistics

| | Mean (deg.) | mean SD (deg.) | mean resultant length | Watson's $U^2$ vs contralateral | Watson's $U^2$ vs DMSO combined |
| --- | --- | --- | --- | --- | --- |
| DMSO control (n = 6) | 226 | 125 | 0.091 | $p > 0.10$ | $p > 0.10$ |
| DMSO contralateral | 213 | 126 | 0.085 | | $p > 0.10$ |
| MEK1/2 inhibitor U0126 (n = 5) | 191 | 127 | 0.025 | $p > 0.10$ | **$p < 0.001$** |
| MEK1/2 inhibitor contralateral | 204 | 136 | 0.048 |  | **$p < 0.001$** |
| PI3K inhibitor LY294002 (n = 5) | 207 | 133 | 0.056 | $p < 0.1$ | **$p < 0.01$** |
| PI3K inhibitor contralateral | 225 | 152 | 0.022 |  | **$p < 0.001$** |
| PLCγ inhibitor U73122 (n = 4) | 230 | 137 | 0.021 | $p < 0.1$ | **$p < 0.001$** |
| PLCγ inhibitor contralateral | 169 | 134 | 0.045 | | **$p < 0.001$** |
| Mixed inhibitor treatment (n = 5) | 212 | 120 | 0.113 | **$p < 0.001$** | **$p < 0.05$** |
| Mixed inhibitor contralateral | 232 | 128 | 0.087 | | $p < 0.1$ |

Golgi positional orientation, or where along the tissue surface the Golgi body is pointing toward, was more ordered than Golgi angle (Figure 7). In all treatment-side combinations the Golgi body tended to lie between the nucleus and the globular process or lateral edge of the FNP on the ipsilateral side. No inhibitor treatments significantly reduced the percentage of cells pointing towards the lateral edge of the FNP but inhibiting MEK1/2 and PLCγ increased the number of cells pointing towards the interior of the head on the ipsilateral side (p = 0.0008 for both, Figure 7). The mixed inhibitor treatment did not affect Golgi positional orientation. Similarly, there were no significant differences between treated and contralateral sides.

**Figure 7:**
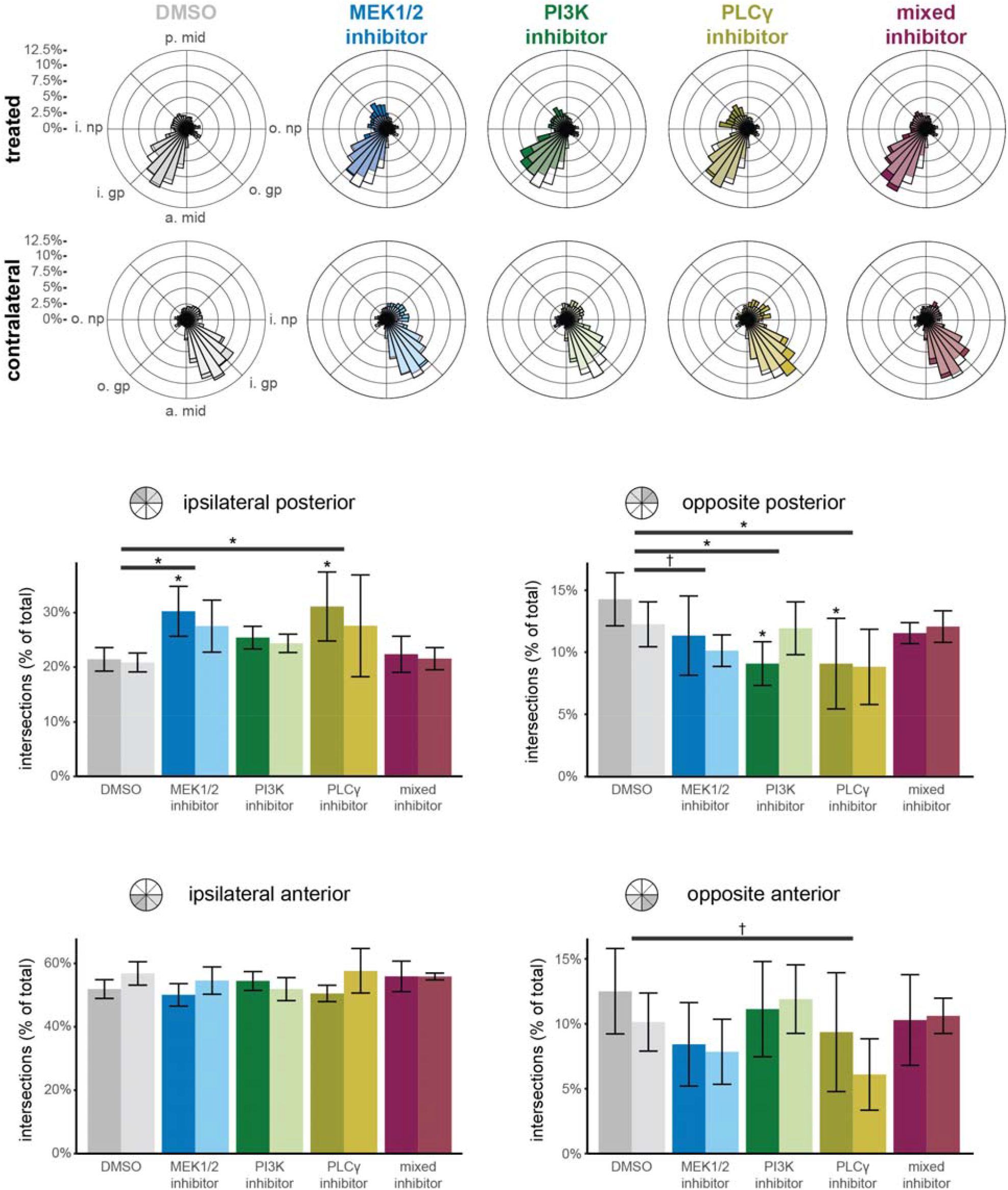
All individual inhibitor treatments, but not the mixed inhibitor treatment, disrupted Golgi positional orientation compared to DMSO control. Positional orientation is illustrated with windrose histograms of percent of nuclei for each treatment and side combination. The ‘angle’ was calculated as the position along the head the Golgi is pointing at. Each windrose histogram shows Golgi positional orientation for the treatment-side combination in color as well as the DMSO combined sides distribution in white. The bottom bar charts show the relative amount of cells pointing at four quadrants of the head. Individual inhibitor treatments, but not the mixed inhibitor treatment, increased the amount of cells oriented away from the ipsilateral anterior portion of the face (nasal pit, globular process, and front of FNP). * p < 0.05, dagger < 0.1, Note: all significant comparisons are between inhibitor vs DMSO or inhibitor-side vs DMSO-side – there were no significant treated vs contralateral side comparisons within treatments.

### Small molecule inhibitors specifically decreased activity of each target pathway

We confirmed the specificity of the small molecule inhibitors using immunohistochemistry to visualize phosphorylated forms of ERK1/2, Akt, and PLCγ in treated and contralateral sides of the FNP six hours after the beads were placed (approximately stage HH20/21). Each molecule decreased phosphorylation of its target protein but not the proteins of other pathways (Figures 8-10).

**Figure 8:**
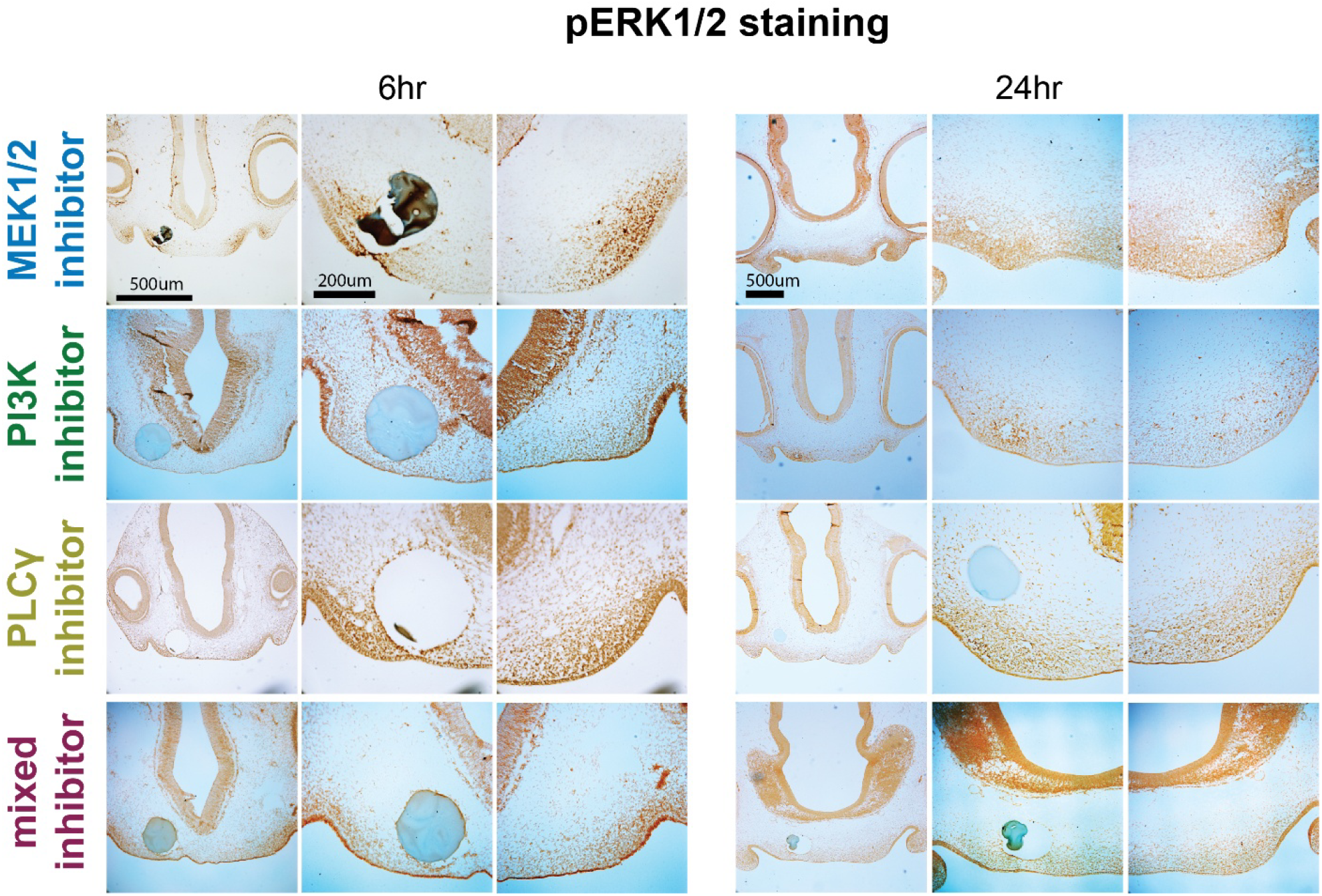
Phosphorylation of ERK1/2. MEK1/2 inhibitor (top row) and the mixed inhibitor (bottom row) decreased the amount of phosphorylated ERK1/2 (pERK1/2) on the bead treated side compared to the contralateral side 6hrs (left columns) and 24hrs (right columns) after treatment. PI3K inhibitor and PLCγ inhibitor treatment did not affect the amount of pERK1/2.

**Figure 9:**
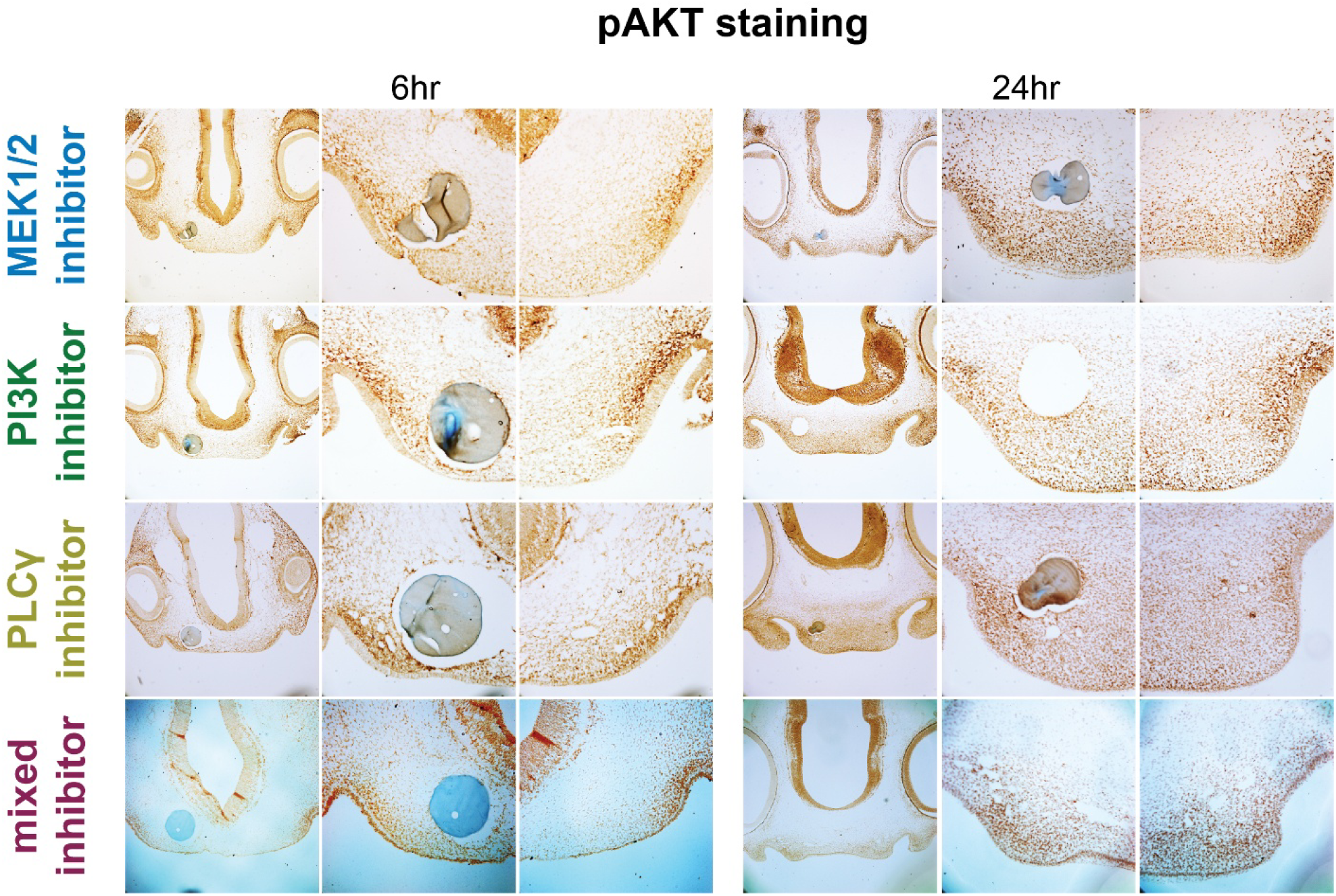
Phosphorylation of Akt. PI3K inhibitor (second row) and the mixed inhibitor (bottom row) decreased the amount of phosphorylated Akt (pAkt) on the bead treated side compared to the contralateral side 6hrs (left columns) and 24hrs (right columns) after treatment. MEK1/2 inhibitor and PLCγ inhibitor treatment did not affect the amount of pAkt. Scale is the same as Figure 8.

**Figure 10:**
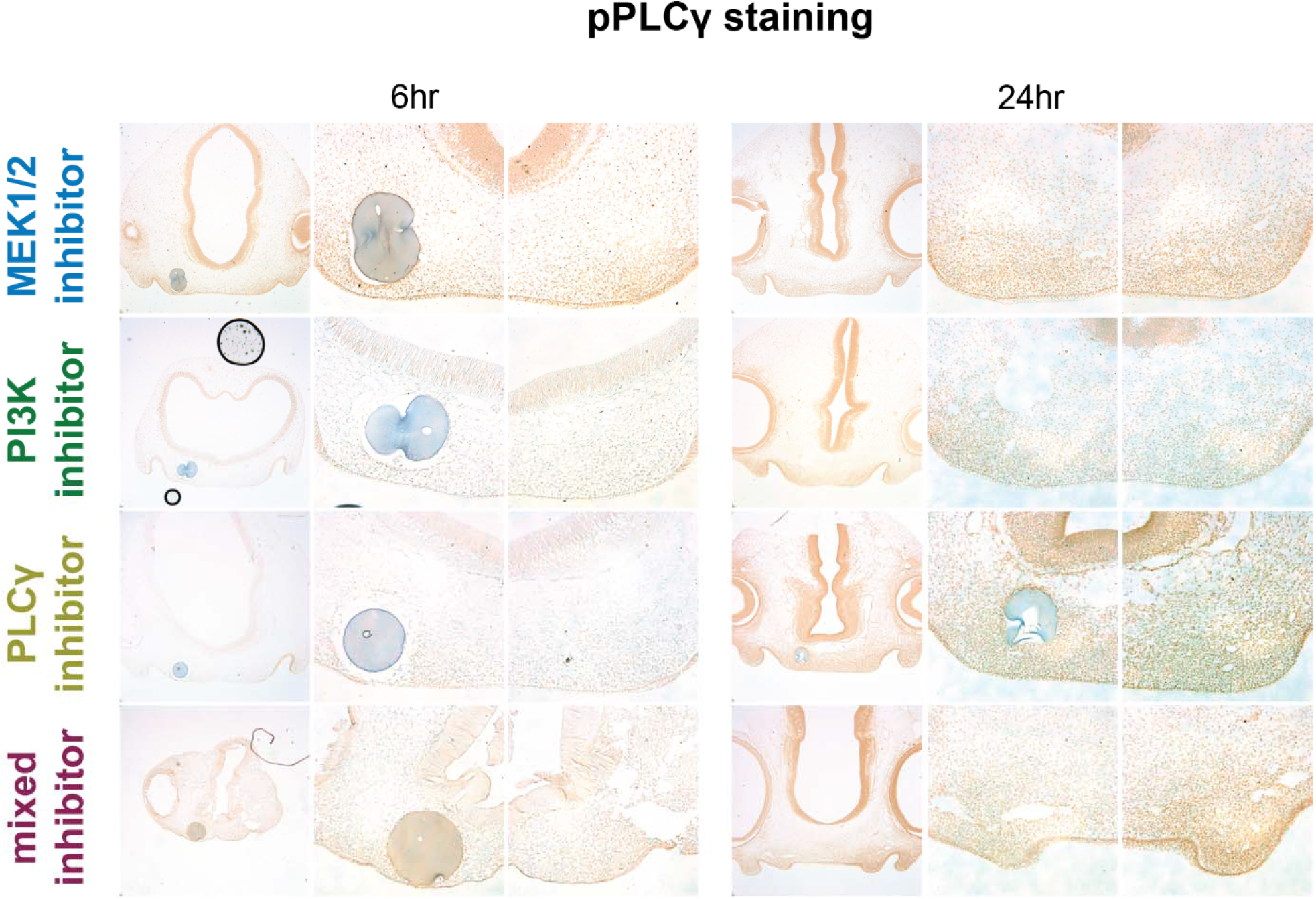
Phosphorylation of PLCγ1. PLCγ inhibitor (third row) and the mixed inhibitor (bottom row) decreased the amount of phosphorylated PLCγ (pPLCγ) on the bead treated side compared to the contralateral side 6hrs (left columns) and 24hrs (right columns) after treatment. MEK1/2 inhibitor and PI3K inhibitor treatment did not affect the amount of pPLCγ. Scale is the same as Figure 8.

## Discussion

Inhibiting three prominent downstream effectors of RTK signaling, MEK1/2, PI3K, and PLCγ caused similar truncation of growth in the frontonasal process. Inhibiting all three downstream pathways simultaneously with the mixed inhibitor treatment caused a slightly more severe truncation of growth, but not as severe as we have observed from inhibiting FGFR signaling directly with SU5402.^21^ Each of the inhibitor treatments caused similar asymmetric truncation on the treated side of the FNP but the effect on shape was highly variable compared to DMSO control and caused symmetric changes on the contralateral, untreated side of the FNP. Although the signaling cascades of MEK1/2, PI3K, and PLCγ are complex, they ultimately affect morphogenesis by changing a small number of cellular activities such as proliferation, apoptosis, orientation of movement and cellular division, cellular shape change, and extracellular matrix production.^25^ Our goal was to determine how each of the downstream pathways of RTK activation affect morphogenesis through changes to cellular activities but found that each inhibitor had similar effects on proliferation, apoptosis, and cellular orientation. Since each of the inhibitors had similar effects on cellular activity and shape of the FNP, and since the mixed inhibitor treatment only slightly increased truncation, the downstream branches of RTK signaling likely have shared, inter-dependent effects on the cellular processes driving morphogenesis.

The mild effect on shape compared to stimulating the FGF pathway with retroviruses or blocking FGFR1 signaling with SU5402 further suggests that robust regulation of RTK signaling compensate for decreased activity of the three suppressed pathways.^10,11,21^ FGFR and PDGFR signaling are required for normal craniofacial development – alteration in RTK signaling results in a wide variety of dysmorphologies and can be embryonic lethal. Alteration in conserved effector pathways that can generate extreme phenotypic outcomes during development are dampened through redundancy, overlapping regulatory networks,^16^ and positive and negative feedback loops.^26–29^ These complex regulatory networks may have contributed to the increased variance in FNP shape observed in the inhibitor treated FNPs. In addition, there are many other RTK functions we did not target with these three inhibitors, including other downstream effector cascades such as c-Jun N-terminal kinase (JNK) and signal transducer and activator of transcription (STAT), crosstalk interactions with other signaling systems such as wnt and Notch, and non-canonical functions of the receptor in cell-matrix interactions.^23,30^ Inhibiting one effector cascade downstream from RTK activation likely affects activation of a different downstream pathway (or another signaling system entirely) to modulate the effect, as has been shown with ERK1/2 activation in response to inhibiting PI3K with LY294002.^16^ The lack of distinct effects of each inhibitor on cellular activity outcomes may be due to redundancy and robust regulatory networks.

Biochemical regulatory network compensation may also explain why the mixed inhibitor treatment did not have as strong an effect as blocking FGFR signaling entirely with SU5402, upstream from all effectors of FGFR activation. However, the mixed inhibitor treatment may have been weaker than individual inhibitors because the concentration of each inhibitor is lower. The beads are made by soaking them in a solution containing 10 mM of all three inhibitors, the highest soluble concentration, potentially diluting each inhibitor by a third compared to the individual treatments. Bead delivery of inhibitors provides transient and spatially limited decrease in kinase activity which could be easier for regulatory networks to compensate for, potentially dampening the full effect of blocking or knocking out downstream pathways. A mouse allelic series of conditional mutations on FGFR1 and FGFR2 proteins’ binding sites that disrupt ERK1/2 and PLCγ signaling caused additive effects when combined, but did not recapitulate the FGFR1/FGFR2 null phenotypes.^23,31^

Inhibiting MEK1/2, PI3K, and PLCγ individually and in tandem on only the right side of the FNP caused symmetric and asymmetric shape change but did not alter covariation of shape between the treated and contralateral sides. The inhibitors altered morphogenesis on both the treated and contralateral sides of the FNP and tended to affect proliferation and Golgi orientation on the contralateral side as well despite the one-sided effect on pathway activation (Figures 8-10). Since there is no known mechanism for coordinating symmetry in the face, the covariation of shape in the contralateral side are potentially caused by biochemical or physical changes. As discussed previously, the effect of the inhibitors is likely dampened through regulatory networks, but it is possible that the boundary of diffusion of the inhibitor and the activity of the networks does not overlap. This could create a gradient of RTK effector signaling within the relatively small volume of the FNP. Changes to cells’ physical environment could alter the forces experienced on the contralateral side of the FNP. Cells create their extracellular matrix which they then use to move, exert force, and sense forces; changes to the ECM due to the inhibitors on the treated side, or changes to pressure due to truncation on the treated side, may alter cellular activity on the contralateral side as they are physically attached to one another.^32^ The hypotheses that biochemical or biomechanical changes contribute to symmetry during morphogenesis warrant further investigation.

Each of the inhibitors reduced cellular proliferation, though inhibiting MEK1/2 with U0126 appeared to have a stronger effect than the other inhibitors. These results are consistent with literature suggesting that all three downstream branches promote proliferation, with MEK1/2 signaling being more associated with FGF signaling and craniofacial development.^14,15,30,33–35^ In our previous work, increasing FGF signaling in the chicken embryo with an RCAS virus expressing *Ffg8* also decreased cellular proliferation, and the decreased proliferation was correlated with proximodistal truncation of the FNP and hypoplasia of the maxillary process.^11^ However, stimulating FGF signaling with the RCAS::*Fgf8* virus increased expression of FGF pathway inhibitors *Spry1*, *Spry2*, *Spred2*, and *Mkp3*/*Dusp6* and decreased the amount of phosphorylated ERK1/2 but not the amount of phosphorylated PLCγ or PI3K, suggesting the decreased proliferation was driven by inhibition of the MEK1/2 pathway.^11^ In this experiment, inhibiting MEK1/2 with U0126 had a relatively mild effect on FNP shape compared to the other treatments. We did not observe increased apoptosis in the FNP with any of our inhibitor treatments. The inhibitor concentrations used in this experiment (10 mM of each inhibitor) are the highest concentrations that can dissolve into solution, but it may be that delivering more inhibitors using multiple beads could cause apoptosis. The strong effect of SU5402 observed in other studies may be due in part to apoptosis as it is cytotoxic.^10,21^ The Soriano group’s FGFR genetic double null conditional knockout (*Fgfr1^cKO/cKO^;Fgfr2^cKO/cKO^)* mouse had increased cell death in the FNP but no effect on proliferation.^23^ Adding a null mutant allele of *Bim* (*Bim^+/−^*), a protein that initiates apoptosis, reduced apoptosis and partially rescued the FNP dysmorphology phenotype in the FGFR double null mice.^23^ PI3K and ERK1/2 activity antagonizes BIM, but so does JNK, which we did not target with any of our inhibitors.^23,31,36,37^ Another study found that inhibiting MEK1/2 and PI3K with small molecule inhibitors increased activation of JNK.^16^

Cellular movement in the direction of growth likely contributes to morphogenesis during development.^6,18^ Previously we have shown that the severity of dysmorphology caused by increasing FGF signaling with retroviruses was associated with decreased consistency of Golgi orientation in the lateral FNP.^11^ Golgi in the DMSO control group tended to orient anterolaterally, towards the globular process, while each of the inhibitor treatments disrupted orientation of the Golgi body with no apparent mean orientation. Surprisingly the mixed inhibitor treated FNPs’ Golgi orientation distributions more closely resembled the DMSO control distribution than the individual inhibitor treated FNPs. One potential reason the mixed inhibitor treatment did not have as strong an effect as the individual inhibitors may be that the beads do not hold as much of each inhibitor. As discussed previously, the mixed treatment has a lower concentration of each individual inhibitor, which could explain the mild effect on Golgi orientation.

We inhibited three signaling cascades downstream from RTK activation to determine the role of each pathway on cellular activities that control tissue growth. We found that each of the pathways had similar effects on cellular activities and on shape change in the FNP and that inhibiting all three pathways together caused a slightly stronger truncation of the FNP but similar changes to cellular activity. We conclude that RTK signaling, a tightly regulated system that is required for growth and formation of all parts of the developing embryo, is robust to developmental perturbation through redundant signaling systems responsible for controlling cellular activities. We did not observe differences in cellular outcomes, but there are other cellular outcomes that were not examined in this work which could have contributed to shape change such as the biomechanical environment of the FNP. Future work will examine the role of biomechanical properties within the FNP on morphogenesis, and how RTK signaling contribute to these properties.

## Experimental Procedures

### Bead implantation

Fertilized chicken eggs (*Gallus gallus*, Rhode Island Red, Petaluma Farms, CA) were incubated in a humidified chamber at 39°C until stage HH19/20.^38^ Ion exchange beads (AG1-X8, 100-200 mesh, 106-180 µm diameter; BioRad, Hercules, CA) were soaked in 10 mM U0126 (9903S, Cell Signaling Technology, Danvers, MA) in dimethylsulfoxide (DMSO), 10 mM U73122 in DMSO (HY-13419, MedChemExpress, Monmouth Junction NJ), 10mM LY294002 in DMSO (A8250 from APExBT, Houston, TX), 10 mM each of U0126, U73122, and LY294002 in DMSO (inhibitor mix), or in DMSO alone as the control. Beads were positioned beneath the ectoderm in the lateral edge of the right globular process of the FNP. Embryos were returned to the incubator for 6 hours (stage HH21/22), 24 hours (stage HH24/25), or 72 hours (stage HH29/30).

### Geometric morphometrics

All geometric morphometric analyses were performed 24 hours after bead implantation when embryos were stage HH24/25. For 3D geometric morphometrics the heads of the embryos were collected, fixed in 4% paraformaldehyde, dehydrated to 70% ethanol, and scanned with microcomputed tomography. Heads were re-hydrated and soaked in a 3.75% iodine solution overnight with gentle shaking the day before scanning. Heads were scanned one at a time in air with a Scanco μCT35, (SCANCO Medical AG, Brüttisellen, Switzerland) using a 12 μm voxel size. 16 fiducial landmarks, 17 curve semi-landmarks, and 18 surface semi-landmarks were placed across reconstructions of the frontonasal process using 3D Slicer (version 4.11) (Figure 1B).^39^ Generalized Procrustes analysis of 3D landmarks were performed using R (version 4.1.2) with the ‘geomorph’ and ‘morpho’ packages.^40,41^

### pHH3 and TUNEL labeling and analysis

Proliferation and apoptosis were analyzed 24 hours after bead implantation when embryos were stage HH24/25. Adjacent transverse sections of the head, immediately cranial to the bead location, were labelled with anti-pHH3 (1:200, 9701S, Cell Signaling Technology, Danvers, MA) or TUNEL *in situ* Cell Death Detection Kit (11684795910, Roche Diagnostics GmbH, Mannheim, Germany) to reveal proliferating and apoptotic cells, respectively. Proliferation was quantified as the ratio of pHH3 positive cells to total nuclei labelled by Hoechst. Digital masks of nuclei and pHH3 labelled cells were created with Cellpose 2.0 and quantified using Python.^42^ A small set of confocal images of nuclei in the same tissue region were traced using the Freehand selection tool in ImageJ to create training data to build a segmentation model in Cellpose. The ‘human-in-the-loop’ training method in Cellpose was used to refine the initial training set with the Hoechst and pHH3 images.^42^ Few TUNEL positive cells were noted so apoptosis was not quantified.

### Golgi body orientation

Golgi body orientation were quantified 24 hours after bead implantation when embryos were stage HH24/25. Angle of the Golgi body relative to the nucleus was measured in three dimensions using confocal images of the FNP (Figure 1C). The nuclei and Golgi bodies were labelled with Hoechst and anti-GM130 (1:500 610823, BD Biosciences, Franklin Lakes, NJ), respectively, on 8-12 μm thick transverse sections of the FNP adjacent to or containing the bead. Stacks with a z-space of 0.4 μm were captured with a confocal microscope (Stellaris 5, Leica Microsystems GmbH, Wetzlar, Germany). The previous segmentation model trained on Hoechst labelled images was refined in Cellpose to create models for nuclei and GM130 labeled images.^42^ Digital masks of nuclei and Golgi bodies were created with Cellpose, and the angles between nuclei masks and Golgi masks were calculated in Python.^42^ Matches were determined using a linear assignment problem with Euclidian distance between centroids as the ‘cost’. Golgi-nuclei pairs were excluded from analysis if they were in the ectoderm, in the neural ectoderm, more than 200 μm away from the ectoderm, or had Golgi separated by more than 25 μm. Since there was little variation in the z-direction, all analysis was performed on 2D angles by removing the z-components. The mean angle, standard deviation, and mean resultant length (1-variance) were calculated in R with the ‘circular’ package.^43^

Golgi positional orientation was calculated using the same digital masks and angles from the Golgi angle analysis described above. Positional orientation was determined by extending the line from the centroid of a nucleus to its’ matching Golgi until it intersected with the edge of the tissue section, and this intersection point was then converted into an angle. The edge of the tissue was composed of eight lines drawn on a low-magnification overview image of each tissue section using the ImageJ freehand line tool. The eight sections (‘octiles’) are: from the midline of the FNP to the globular process, from the globular process to the top of the nasal pit, from the top of the nasal pit to neural ectoderm roughly in-line with the globular process, from that neural ectoderm to the neural ectoderm midline, and then these same four regions on the opposite side of the head. The octiles were divided equally across a circle, so each quadrant was 45° apart with 0° and 180° centered on the top of the nasal pits and 90° and 270° on the neural and FNP midline points, respectively (Figure 1C). Positional angle was calculated as a ratio of the length along an octile to the point where the Golgi line intersects the octile, and the total length of the octile. For example, if a line from a nucleus through its’ Golgi body then intersects the tissue just lateral of the FNP midline, the angle would be 270° + (length from midline to intersection / length from midline to globular process) x 45°.

### Immunohistochemistry

Immunohistochemistry for phosphorylated forms of ERK1/2, Akt, and PLCγ were performed on paraffin-embedded sections of samples harvested 6hr and 24hr after beads were implanted (n > 5 per antibody). Antibodies specific to phosphorylation on ERK1/2 threonine 202 and tyrosine 204 were used to detect pERK1/2 (1:100, 4370S, Cell Signaling Technology, Danvers, MA); antibodies specific to phosphorylation of serine 473 of Akt1/2/3 were used to detect pAkt (1:300, 4060S, Cell Signaling Technology, Danvers, MA); and antibodies specific to phosphorylation of tyrosine 783 of PLCγ1 were used to label pPLCγ (1:250, BS-3343R, Bioss, Woburn, MA). The basic protocol included antigen retrieval in 10 mM tris-EDTA buffer (10 min, 100°C), endogenous peroxidase blocking in 3% H_2_O_2_ (10 min, room temperature) and nonspecific epitope blocking with 5% goat serum (60 min, room temperature). Primary antibodies were applied to sections overnight at 4LC. A horseradish peroxidase-conjugated, species-specific secondary antibody (1:200 in phosphate-buffered saline with 5% goat serum, AB307P, MilliporeSigma, Burlington, MA) was incubated for 60 min at room temperature, then detected using peroxidase substrate kit (SK-4100, Vector Laboratories, Newark, CA).

The sections were cover-slipped before imaging on a Leica DMRB brightfield microscope (Leica Microsystems GmbH, Wetzlar, Germany). Brightness, contrast, and white balance were adjusted using Photoshop 2025 (Adobe, San Francisco, CA).

### Statistical analysis

All statistical analyses were performed using R to answer the following questions:

1. What is the effect of each treatment on mean shape and variation in shape of the FNP compared to the DMSO control treatment? Was the shape change caused by treatments symmetrical or asymmetrical?
2. What is the effect of each treatment on cellular proliferation compared to the DMSO control treatment? What is the effect of each treatment on cellular proliferation between the treated and contralateral side of the FNP?
3. What is the effect of each treatment on the distribution of Golgi orientation between treated and contralateral sides of the FNP? What is the effect of each treatment compared to the DMSO control treatment?
4. Generalized Procrustes analysis (GPA) was performed on 3D landmark data and shape was visualized using principal components analysis (PCA). We quantified the relationship between shape of the FNP to size of the FNP (centroid size) with a general linear model and found a significant effect on allometry between DMSO and inhibitor treated samples. As described in the results, we removed the effect of allometry by subtracting the regression of the residuals of the first principal component from the original landmark coordinates. All subsequent analysis of shape was performed with these transformed coordinates. The effect of treatment on mean shape (GPA transformed landmark coordinates) of the FNP was examined using a general linear model with pairwise comparisons between different treatment groups. The degree of symmetry and asymmetry were analyzed in the same way as mean shape, but the GPA was performed with the assumption that the landmark data exhibited bilateral symmetry about the midline of the FNP (Fig 1B). The symmetric and asymmetric components were compared using a general linear model and pairwise comparisons. The ‘geomorph’ package was used to perform GPA and solve the linear model, while the ‘pairwise’ package was used to perform treatment group comparisons. Heatmaps comparing shape at different principal component scores were generated using the ‘morpho’ package. Covariation of shape between the inhibitor treated side and the contralateral, untreated side were compared using the ‘integration.test’ in the ‘geomorph’ package. We generated a paired, two-block partial least squares analysis (PLS) between the two sides of the FNP and the effect size of each inhibitor PLS was compared to the DMSO PLS using the ‘compare.pls’ function, which computes a two-sample z-test of the standard deviations of each PLS.^40^ We compared faces composed of mirrored halves to further examine the effect of treatment on the contralateral, untreated side (Fig 1B). New FNP samples were created by mirroring one side of the FNP across the midline to create ten new groups of samples: DMSO-treated (composed of the landmarks from the right, DMSO bead treated side of the FNP, the midline landmarks, and the right side mirrored over to the left, contralateral side), DMSO-contralateral, mixed-treated, mixed-contralateral, etc. The same GPA and PCA analyses described above were used to analyze shape in these new samples.
5. Cellular proliferation was compared between the bead treated and contralateral sides of the face using a paired Student’s t-test for each treatment group. Proliferation in the contralateral side of U0126 and LY294002 groups were compared to the combined treated and contralateral sides of the PLCγ group using a Welch’s two-sided t-test.
6. Golgi angles were simplified to only two-dimensions by removing the z-component. All angle measurements on the contralateral side of the FNP were mirrored across the ‘yz’ plane to align with the bead treated side. The distribution of 2D angles were compared between treatment groups and treated or contralateral sides of the FNP using Watson’s non-parametric two sample U^2^ test from the R package ‘circular’.^43,44^ This test determines if the distribution of angles of two given samples could have been drawn from the same population. Note, this is not a paired test. First, the bead treated sides were compared to the contralateral sides for each treatment (i.e. U0126 bead treated sides compared to U0126 contralateral sides). Finally, since the DMSO treated and contralateral sides were similar, both DMSO sides were combined into a single distribution and compared to each treatment group and side combination (i.e. DMSO combined sides of the FNP compared to U0126 bead treated sides).

Golgi positional orientation was converted to an angle for visualization with windrose plots. Golgi positional orientation is not true angular data, so analysis with Watson’s test would not appropriate. All positional orientation measurements on the contralateral side of the FNP were mirrored across the ‘yz’ plane to align with the bead treated side. The relative number of intersections in four regions were compared between treatment and side using ANOVA with Tukey’s post-hoc comparisons in R.

## Acknowledgments

We thank all members of the Orthopaedic Trauma Institute Laboratory for Skeletal Regeneration for support throughout this project, especially Nathan Young for his help with morphometrics, and Gina Baldoza for laboratory management. We also thank the staff within the Biological Imaging Development CoLab (BIDC) at UCSF Parnassus Heights for their training and support with confocal microscopy.

